# The gut-brain vagal axis scales hippocampal memory processes and plasticity

**DOI:** 10.1101/2024.04.28.591513

**Authors:** Oriane Onimus, Faustine Arrivet, Isis Nem de Oliveira Souza, Benoit Bertrand, Julien Castel, Serge Luquet, Jean-Pierre Mothet, Nicolas Heck, Giuseppe Gangarossa

## Abstract

The vagus nerve serves as an interoceptive relay between the body and the brain. Despite its well-established role in feeding behaviors, energy metabolism, and cognitive functions, the intricate functional processes linking the vagus nerve to the hippocampus and its contribution to learning and memory dynamics remain still elusive.

Here, we investigated whether and how the gut-brain vagal axis contributes to hippocampal learning and memory processes at behavioral, functional, cellular, and molecular levels. Our results indicate that the integrity of the vagal axis is essential for long-term recognition memories, while sparing other forms of memory. In addition, by combing multi-scale approaches, our findings show that the gut-brain vagal tone exerts a permissive role in scaling intracellular signaling events, gene expressions, hippocampal dendritic spines density as well as functional long-term plasticities (LTD and LTP). These results highlight the critical role of the gut-brain vagal axis in maintaining the spontaneous and homeostatic functions of hippocampal ensembles and in regulating their learning and memory functions.

In conclusion, our study provides comprehensive insights into the multifaceted involvement of the gut-brain vagal axis in shaping time-dependent hippocampal learning and memory dynamics. Understanding the mechanisms underlying this interoceptive body-brain neuronal communication may pave the way for novel therapeutic approaches in conditions associated with cognitive decline, including neurodegenerative disorders.

**Highlights:**

- The gut-brain vagal axis contributes to long-term recognition memories
- The gut-brain vagal axis is dispensable for short-term memories
- The vagal axis regulates molecular and signaling dynamics in the hippocampus
- The gut-brain vagal tone shapes the structural density of hippocampal dendritic spines
- The gut-brain vagal tone ensures physiological forms of synaptic plasticity

## Introduction

The vagus nerve represents the major interoceptive relay allowing the bidirectional neuronal communication between the body and the brain. While its descending efferent component ensures the parasympathetic control of peripheral organs, the ascending afferent component continuously signals to the brain the metabolic and physiological states of several peripheral organs (*i.e.* lung, heart, gut). Vagal afferents-conveyed sensory information is processed and integrated by neurons harbored within the brainstem region of the nucleus of the tractus solitarius (NTS) which in turn projects to a variety of downstream brain regions (Dumont et al., 2022; Gasparini et al., 2020; He et al., 2022; Roman et al., 2016; Trapp and Cork, 2015), therefore contributing to the establishment of extensive interoception-related neuronal networks.

In addition to its crucial role in regulating feeding- and metabolism-associated features (Berthoud and Neuhuber, 2019), the vagus nerve has been pointed as a modulator of cognitive and mnemonic processes. Indeed, electrical stimulation of the vagus nerve facilitates behavioral learning and memory functions (Clark et al., 1999, 1998), promotes hippocampal neurogenesis and plasticity-associated features (Biggio et al., 2009; Follesa et al., 2007). In addition, few but seminal recent articles have also shown that the disruption of the vagal axis, especially at the gut-brain segment, leads to impairments of some forms of hippocampal memory along with cellular and molecular adaptations (O’Leary et al., 2018; Suarez et al., 2018).

Indeed, the hippocampus represents one the main structures involved in the control of learning and memory dynamics. The anatomical and functional links between the vagus nerve and the hippocampus are not direct but require the mobilization of brainstem/hindbrain-reaching intermediary structures which ultimately project to the hippocampus, such as the medial septum (Suarez et al., 2018), locus coeruleus (Kempadoo et al., 2016), hypothalamic and thalamic nuclei (Tao et al., 2021).

Noteworthy, mounting experimental evidence indicates that neurodegenerative disorders including Parkinson’s and Alzheimer’s diseases, which lead to substantial learning and memory alterations, are associated with physiopathological dysfunctions of the gut-brain vagal axis (Singh et al., 2021). In fact, the vagus nerve, at least in part, contributes to the gut-to-brain neuronal seeding and spreading of neurodegenerative hallmarks (*i.e.* misfolded α-synuclein and β-amyloid proteins) (Braak et al., 2003; Chandra et al., 2023; Chen et al., 2021; Kim et al., 2019; Sun et al., 2020), therefore strongly participating in the cognitive decline observed in neurodegenerative disorders. Altogether these breakthroughs are progressively leading to the establishment of less *braincentric* and more integrative/holistic hypotheses of memory-related regulatory elements.

However, while this compelling body of evidence highlights a memory-associated role of the vagal axis, the behavioral, cellular and molecular vagus-dependent underpinnings involved in gating hippocampal learning and memory dynamics remain still elusive.

The present study investigated the role of gut-brain vagal signaling on time-dependent (short- and long-term) hippocampal memories as well as on cellular and molecular (mal)adaptations following perturbations of the gut-brain vagal integrity. Our results indicate that this integrity is necessary for long-term recognition memories, while sparing short-term dynamics. We also demonstrate that the integrity of the gut-brain vagal axis ensures the endogenous/constitutive remodeling of hippocampal dendritic spines as well as the induction of cell signaling events and the transcriptional expression of different genes involved in neuronal transmission, spines’ morphogenesis and plasticity. In addition, we provide evidence that the gut-brain vagal axis exerts a key role in scaling even long-term forms of elicited synaptic plasticity (namely LTP and LTD), thus indicating that interoceptive signals spontaneously participate in the correct framing and maintenance of hippocampal functions.

## Material and methods

### Animals

All experimental procedures were approved by the Animal Care Committee of the Université Paris Cité (CEB-22-2019, APAFiS #24407) and the University Paris-Saclay (accreditation number C92-019–01), and carried out following the 2010/63/EU directive. 8-12 weeks old C57BL/6J male mice (Janvier, France) were used and housed in a room maintained at 22 +/-1 °C, with a light period from 7h00 to 19h00. Regular chow diet (3.24 kcal/g, reference SAFE® A04, Augy, France) and water were provided ad libitum unless otherwise stated.

### Subdiaphragmatic vagotomy

Prior to surgery and during 3 post-surgery days, animals were provided with *ad libitum* jelly food (DietGel Boost #72-04-5022, Clear H2O). Animals received Buprécare® (buprenorphine 0.3 mg/kg) and Ketofen® (ketoprofen 10 mg/kg) and were anaesthetized with isoflurane (3.5% for induction, 1.5% for maintenance). During surgery the body temperature was maintained at 37°C. Briefly, using a binocular microscope, the right and left subdiaphragmatic branches of the vagus nerve were carefully isolated along the lower esophagus/stomach and carefully sectioned in vagotomized animals (SDV) or left intact in sham animals. Mice recovered for at least 3-4 weeks before being used for experimental procedures.

### Novel object recognition (NOR) memory

The novel object recognition (NOR) test was performed as previously described (Gangarossa et al., 2014a, 2014b). Briefly, mice were habituated to a V-maze with two identical arms (34 × 6 × 15 cm, at 90°) for 10 min. The following day, mice were allowed to freely explore two objects located at the end of the two arms. Object interaction was defined as approaching the object with the nose closer than 1 cm. Following a retention interval of 30 min (short-term memory) or 24 h (long-term memory), mice underwent a 5 min recall session during which the V-maze contained a familiar object and a novel object. Following each session, the objects and the maze were cleaned with 70% ethanol. The experiments were videotaped and the time spent exploring the objects was blindly scored by a second experimenter.

### Novel place recognition (NPR) memory

The novel place recognition (NPR) test was performed as previously described (Gangarossa et al., 2014a, 2014b). Mice were habituated to an arena for 10 min. The following day, mice were allowed to freely explore two objects located in the same site of the arena. Object interaction was defined as approaching the object with the nose closer than 1 cm. Following a retention interval of 30 min (short-term memory) or 24 h (long-term memory), mice underwent a 5 min recall session during which the arena contained one of the two objects displaced in the other site of the arena (novel place). Following each session, the objects and the arena were cleaned with 70% ethanol. The experiments were videotaped and the time spent exploring the objects was blindly scored by a second experimenter.

### T-maze working memory

The T-maze apparatus consisted of three arms (1 main alley and 2 identical arms). This test relies upon the innate tendency of mice to explore novel environments. During the familiarization/training period (10 min), the access to one of the two identical arms was blocked through a sliding door and each mouse was allowed to explore the other arm. Following a retention interval of 30 min (short-term memory) or 24 h (long-term memory), mice underwent a 5 min recall session during which the sliding door was removed (new arm) having full access to both the familiar and new arms. The number of entries were used as index of spatial discrimination.

### Spontaneous alternation Y-Maze

The Y-maze apparatus had three identical arms (40 × 9 × 16 cm) placed at 120° with respect to each other. As previously described (Forner-Piquer et al., 2021), each mouse was placed at the end of one arm and allowed to explore freely the apparatus for 5 min. Spontaneous alternation performance (SAP) was assessed by scoring the pattern of entries into each arm during the 5 min of the test. Alternations were defined as successive entries into each of the three arms as on overlapping triplet sets (*i.e.*, 1-2-3, 2-3-1, …). Percent of spontaneous alternations was defined as the ratio of actual (= total alternations) to possible (= total arm entries −2) number of alternations × 100. Total entries were scored as an index of ambulatory activity in the Y-maze.

### Rotarod test

Balance and motor coordination as well as motor learning were assessed using a mouse accelerating rotarod (Ugo Basile, Comerio, Italy). Mice were placed on the rotating drum that accelerated from 4 to 40 rpm over 5 min for three trials a day, for 4 consecutive days. The trial interval was 45 min for all the mice. Rotarod performances were scored for latency to either fall or ride around the rod.

### Exploration and recognition of a non-familiar environment

Mice were exposed during two consecutive days to a non-familiar environment (rectangular arena with fresh bedding) for 120 minutes and then replaced in their home cages. Locomotor activity (beam breaks, bb) was measured in an automated online measurement system using an infrared beam-based activity monitoring system (Phenomaster, TSE Systems GmbH, Bad Homburg, Germany). A reduced exploratory drive during the second exposure indicates habituation and intact environmental recognition.

### Quantitative RT-PCR

The dorsal hippocampus was dissected from coronal brain sections obtained using a stainless-steel matrix with 0.5-mm coronal sections interval and snap-frozen using liquid nitrogen. All tissues were kept at −80°C until RNA extraction. Tissues were homogenized in TRIzol/QIAzol Lysis Reagent (Life Technologies) with 3 mm tungsten carbide beads by using the Tissue Lyser.

Total RNA was extracted using the RNeasy Micro Kit (QIAGEN). The RNA was quantified by using the NanoDrop 1000 spectrophotometer. 1μg of mRNA from each sample was used for retrotranscription, performed with the SuperScript®III Reverse Transcriptase (Life Technologies) following the manufacturer’s instructions.

Quantitative RT-PCRs were performed in a LightCycler 1.5 detection system (Roche, Meylan France) using the Takyon No Rox SYBR MasterMix dTTP Blue (Eurogentec) in 384-well plates according to the manufacturer’s instruction. All primer sequences used in this study are provided in **Suppl. Table 1**. Relative concentrations were extrapolated from the concentration range for each gene. Concentration values were normalized to the house-keeping gene RPL19.

### Tissue preparation and immunofluorescence

Mice were anaesthetized with pentobarbital (500 mg/kg, i.p., Sanofi-Aventis, France) and transcardially perfused with cold (4°C) PFA 4% for 5 minutes. Sections were processed and confocal imaging acquisitions were performed as previously described (Berland et al., 2020; Gangarossa et al., 2019). The following primary antibodies were used: rabbit anti-Ser^235/235^-S6 (1:500, Cell Signaling) and guinea pig anti-cFos (1:500, Synaptic Systems). Quantification of cFos- and phospho-S6-immunopositive cells was performed using the cell counter plugin of ImageJ taking a fixed threshold of fluorescence as standard reference.

### Dendritic spines morphology

Dendrites and dendritic spines were labeled using the Diolistic technique as previously described (Grutzendler et al., 2003; Heck et al., 2012). Briefly, 3 mg of solid red DiI (1,1’-dioctadecyl-3,3,3’,3’-tetramethylindocarbocyanine perchlorate, Molecular Probes) dissolved in methylene chloride were mixed with 50 mg of tungsten beads (1.3 microns in diameter, Bio-Rad). Methylene chloride was allowed to evaporate, so the DiI adhered to the beads. Beads covered with DiI were coated on the inner surface of a Teflon tube, which was pretreated with polyvinylpyrrolidone (10 mg/ml in water, Sigma-Aldrich). The tube was then cut in pieces, which were inserted as cartridges in a gene gun device (Bio-Rad). Helium gas pressure (150 psi) applied through the gene gun ejected the beads out of the cartridge onto the brain section. The beads were delivered through a 3-µm pore-size filter (Isopore polycarbonate, Millipore) to avoid clusters. A 2 hours incubation in PBS allowed passive diffusion of the DiI in plasma membranes of neurons before the sections were mounted in Prolong Gold.

Images stacks were taken with a Confocal Laser Scanning Microscope (SP5, Leica) equipped with a 1.4 NA objective (oil immersion, Leica) with pinhole aperture set to 1 airy unit, pixel size of 60 nm and z-step of 200 nm. The excitation wavelength and emission range were 561 and 570–650 nm. Laser intensity was set so that each image occupied the full dynamic range of the detector, a low noise Hybrid detector combining avalanche gain and GaAsP detection (HyD Leica).

Deconvolution with experimental point spread function obtained from 100 nm fluorescent beads was performed with Huygens software (Scientific Volume Imaging). One hundred fifty iterations were applied in classical mode, background intensity was averaged from the voxels with lowest intensity, and signal to noise ratio values were set to a value of 20.

Neuronstudio software (version 0.9.92) (Rodriguez et al., 2008) was used to reconstruct the dendrite and detect dendritic spines. Manual correction was required for a minority of spines. For the analysis of dendrite diameter, dendrite models obtained from Neuronstudio and the average diameter of each dendrite was computed with the software L-Measure (Scorcioni et al., 2008). Stubby spines are defined as spines without a neck. While head diameter to neck diameter ratio usually arbitrarily defines thin/mushroom categories, Neuronstudio failed to detect necks in our image stacks (when the spine neck is not detected, Neuronstudio retrieves the tenth of the xy voxel size as default neck diameter), then thin and mushroom spines are categorized by Neuronstudio as follows: Neuronstudio computes the size of the spine head in 2D in the xy plane of the image stack and tag as mushroom all spines having a head diameter wider that 350 nm. Frequency distribution of spine head volumes (μm^3^) for spines with necks were analyzed after 3D segmentation with custom procedure. For this, spine head volumes were extracted as follows. Data generated by Neuronstudio are saved in text file format, which is imported into ImageJ. The head spine center of mass was read in the Neuronstudio table and the closest local maximum was found in the image stack. Each local maximum thus marked a spine head which was then segmented using our spot segmentation procedure (Heck et al., 2015). The 3D intensity distribution around the local maximum was computed and fitted with a Gaussian curve to determine the intensity of the edge of the spine head. Starting from the local maximum, voxels are iteratively added when meeting two intensity criteria: being above the threshold defined for the edge, and being lower than the voxels previously added. The second criterium has been set to prevent the merge in one object of two adjoining spines. Length of the spine was an approximate by measuring the 3D distance between the geometrical center of the spine head segmented with our method and the surface of the vertices from the dendrite model reconstructed in Neuronstudio and imported in ImageJ. For CA1 hippocampal neurons, secondary apical dendrites were analyzed. For DG neurons, dendrites in the molecular layer were analyzed. Each dendrite segment analyzed was 40 to 80 μm in length, with each dendrite belonging to another neuron.

### *Ex vivo* electrophysiology

#### Slice preparation

Animals were deeply anesthetized with isoflurane before decapitation. The brain was then quickly removed and placed in cold (0-4°C) artificial cerebrospinal fluid (aCSF) containing (in mM): NaCl 124, KCl 3.5, MgSO_4_ 1.5, CaCl_2_ 2.5, D-glucose 11, NaH_2_PO_4_ 1.2, NaHCO_3_ 26.2 (pH 7.3-7.4 and 302-313 mOsml/L), previously and continuously bubbled with 95 %O_2_/5 % CO_2_. 400 μm thick coronal slices of the hippocampus were obtained using a 7000smz-2 vibratome (Campden Instruments, USA) and allowed to recover in aCSF for 1 hour at 26°C. After that time, slices were maintained at room temperature (21-26°C) for the remainder of the day in bubbled aCSF. Recordings were performed between 1h 30min and 6h after decapitation.

#### Electrophysiology and synaptic plasticity

Slices were individually transferred to a recording chamber mounted on an upright Slicescope Pro 6000 system (Scientifica, UK) and continuously superfused with oxygenated aCSF (2-3 mL/min) at 28-30°C (Heated perfusion tube HPT-2A, ALA Scientific Instruments Inc, USA). A glass micropipette filled with 2M NaCl (2–4.5MΩ) was placed 150-200 μm deep into the CA1 stratum radiatum for extracellular field potential recording, and a bipolar insulated tungsten microelectrode (FHC, USA) was placed at the Schaffer collaterals fibers for stimulation, 200-350μm apart from the recording electrode. Recordings were made using a Multiclamp 700B amplifier (Molecular devices, CA, USA) and signals were obtained in bridge mode, amplified 200x, filtered at 3 kHz and digitized at 10 kHz via a Digidata 1440A interface (Molecular Devices, CA, USA). Experimental protocol was adjusted from (Le Bail et al., 2015): an input-output curve was obtained to choose a stimulation intensity sufficient for ~0.1 mV/ms response slope for long-term potentiation (LTP) experiments and ~0.15 mV/ms for long-term depression (LTD) experiments (14.5-35 µA, 100 μs, DS3 Constant Current Stimulator, Digitimer, USA). After stimulation intensity was chosen, in LTP experiments, a paired-pulse protocol consisting of two single pulses 40 ms apart was done (2 sweeps, 15 s interval) to defined pre-conditioning paired-pulse facilitation. Immediately after, baseline acquisition consisted of single pulses at the chosen stimulation intensity at 15 s intervals for 10 min. After baseline acquisition, one of two conditioning protocols was used: LTP was induced using a high frequency stimulation (HFS) protocol consisting of 2 trains of pulses at 100 Hz for 1 s with 10 s interval between trains. LTD was induced using paired-pulse low frequency stimulation (PP-LFS) protocol consisting of two pulses with 40 ms interval at 1 Hz for 15 min (Kemp and Bashir, 2001). Immediately after conditioning, single pulse stimulation resumed for 60 min. After 60 min, in LTP experiments only, paired-pulse protocol was repeated to define post-conditioning paired-pulse facilitation (4 sweeps, 15 s interval).

#### Analysis

Data were analyzed using pClamp10 software (Molecular Devices, CA, USA). For paired-pulse ratio analysis, the average of 2 to 4 sweeps were used to measure the 20-80% slope of the two fEPSPs. After, the value of the second pulse is divided by the first to obtain the ratio. For LTP/LTD analysis, the average of 4 sweeps (1 min of recording) was used to measure the 20-80% slope of fEPSPs. The mean value of baseline recordings was considered 100% and the response post-conditioning is plotted as % of that value. Finally, the mean value of the last 10 minutes of recording was obtained and unpaired Student’s t-tests performed to evaluate differences between groups. For PP-LFS protocol analysis, the average of 60 sweeps (1 min of recording) is presented as percentage of baseline. Experiments with 1) more than 15% variation from mean on any baseline value or 2) mean of last 10 min of potentiation/depression below 3 standard deviations for each slice were excluded from analysis and considered unsuccessful.

### Statistics

All data are presented as mean ± SEM. Statistical tests were performed with Prism 7 (GraphPad Software, La Jolla, CA, USA). Detailed statistical analyses are listed in the **Suppl. Table 2**. Depending on the experimental design, data were analyzed using either Student t-test (paired or unpaired) with equal variances, one-way ANOVA or two-way ANOVA. The significance threshold was automatically set at p<0.05. ANOVA analyses were followed by Bonferroni post hoc test for comparisons only when overall ANOVA revealed a significant difference (at least p<0.05).

## Results

### Long-term recognition memories depend on the integrity of the gut-brain vagal axis

The vagus nerve has recently been pointed as a contributor of learning and memory processes (Olsen et al., 2023). By taking advantage of the subdiaphragmatic vagotomy model (SDV), here we investigated whether the gut-brain vagal axis plays a permissive role in shaping learning and memory events, notably recognition memories which strongly rely on the hippocampus formation (Broadbent et al., 2004; Gálvez-Márquez et al., 2022).

First, sham and SDV mice underwent a novel object recognition test (NOR, **Fig. 1A**) which assesses the capacity of an animal to discriminate between a familiar and a novel object. As expected, during the recall session (24 hours after the familiarization session), we observed that sham mice spent more time exploring the novel object as compared to the familiar one (**Fig. 1B**), therefore indicating intact memory recall functions. On the contrary, SDV mice showed an equivalent exploration time between the two objects (**Fig. 1B**), revealing that the integrity of the gut-brain vagal axis is essential for encoding memory-associated long-term events. This was further highlighted by the discrimination index which showed that sham mice had higher performances than SDV mice (**Fig. 1C**). Importantly, this behavioral difference was not attributed to an imbalanced interaction with objects during the training/familiarization phase (**Fig. 1D**).

**Figure 1.**
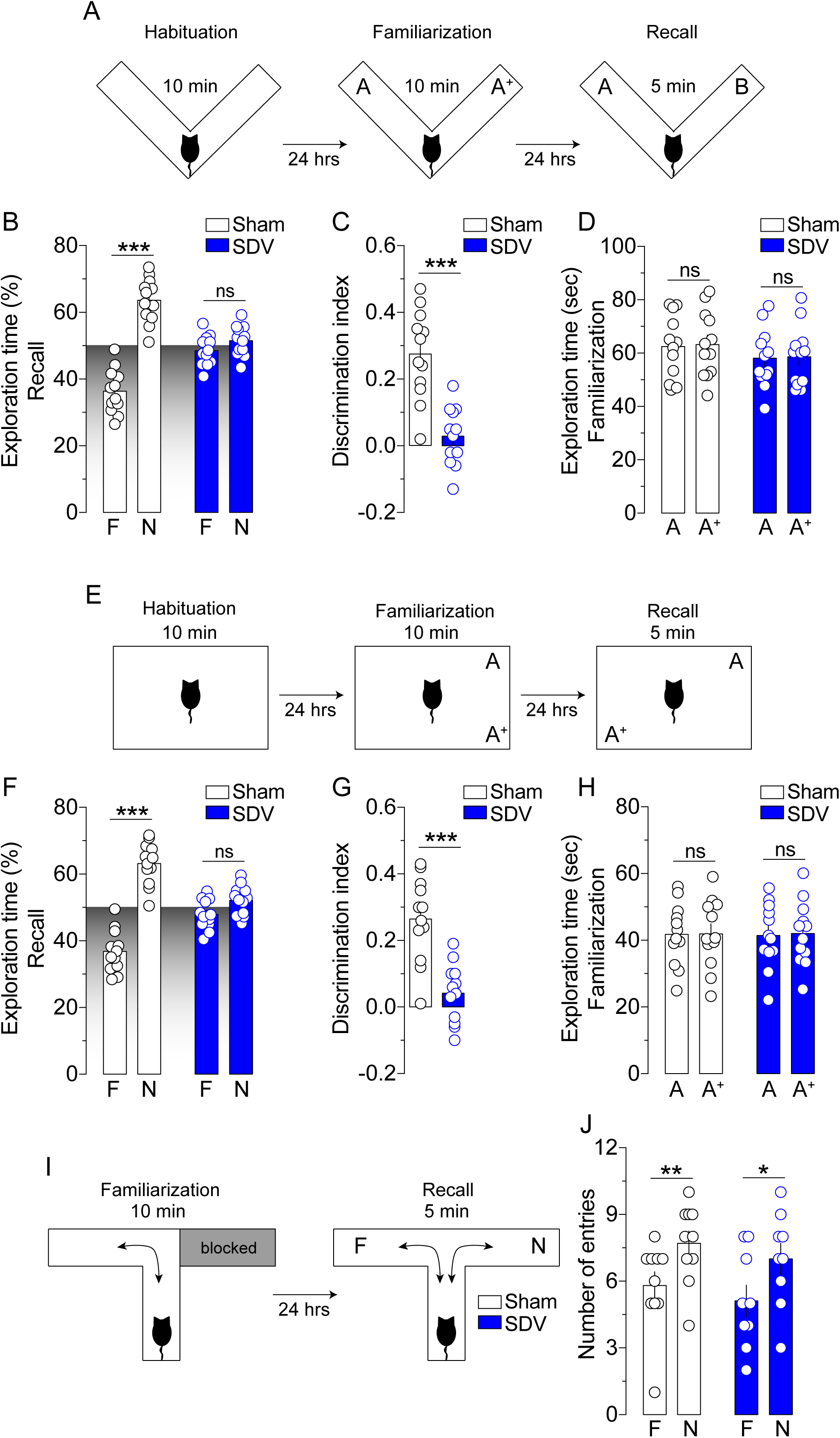
The gut-brain vagal axis regulates long-term recognition memory. (A) The drawing represents the procedure used for the long-term novel object recognition (NOR) test. (B) Percentage of total exploration time for sham and SDV mice exploring the familiar (F) and the novel (N) object. (C) Recognition memory discrimination index. (D) Time spent by sham and SDV mice familiarizing with the two objects. (E) The drawing represents the procedure used for the long-term novel place recognition (NPR) test. (F) Percentage of total exploration time for sham and SDV mice exploring the familiar (F) and the novel (N) places where objects have been displaced. (G) Recognition memory discrimination index. (H) Time spent by sham and SDV mice familiarizing with the two objects. (I) The drawing represents the procedure used to test long-term reference spatial memory by the T-Maze test. (J) Number of entries in the familiar and new compartment. Statistics: *p<0.05, **p<0.01, ***p<0.001 for Sham^(F)^ *vs* Sham^(N)^ (B, F, J) or SDV^(F)^ *vs* SDV^(N)^ (J). Statistics: ***p<0.001 for Sham *vs* SDV mice (C and G). For number of mice/group and statistical details see **Suppl. Table 2**.

Then, we performed a novel place recognition test (NPR, **Fig. 1E**) which assesses the ability of an animal to discriminate between a familiar and a novel location of two identical objects. Interestingly, we observed that SDV mice, as compared to sham mice, failed in distinguishing between the familiar and novel location (**Fig. 1F, G**), even though both experimental groups similarly explored the two objects during the prior 24 hrs training/familiarization phase (**Fig. 1H**). Altogether, these results show that the vagal axis plays a permissive role in mediating recognition memories.

Since the NPR test relies on both recognition (objects) and recognition/spatial (location) memories, to further extend these results, we investigated whether the integrity of the gut-brain vagal axis would be important for long-term spatial reference memory. To this end, we used the T-maze test which assesses the ability of an animal to discriminate between a familiar and a novel environment (**Fig. 1I**). No differences were observed between the two experimental groups (**Fig. 1J**), indicating that recognition impairments do not depend on proper spatial/environmental dysfunctions.

We then wondered whether the memory-related impairments observed in SDV mice could be generalized to other forms of hippocampus-independent long-term memories (*i.e.* motor learning and memory, habituation to a novel environment). In the rotarod test, which mainly reflects motor learning and memory processes dependent on striatal (Dang et al., 2006; Yin et al., 2009) and cerebellar ensembles (De Zeeuw and Ten Brinke, 2015; Galliano et al., 2013), sham and SDV mice showed intact capacities to learn and improve their motor performances over training days (**Suppl. Fig. 1A**). Of note, both groups showed similar inter-trials motor learning during the training phase (inset of **Suppl. Fig. 1A**). In addition, we also tested the capacity of a mouse to recognize a non-familiar environment (arena) by measuring its locomotor activity during two consecutive explorations over two consecutive days. Sham and SDV mice showed similar locomotor activities [explorations during Day1 (D1) and Day2 (D2)] (**Suppl. Fig. 1B**) as well as similar habituation phases (**Suppl. Fig. 1B^1^**), thus suggesting unchanged exploratory and motor functions, and intact environmental recognition.

### The gut-brain vagal axis is dispensable for short-term recognition memories

Given the above-mentioned results (**Fig. 1**), we decided to investigate whether the vagus nerve was also essential in mediating short-term recognition memories. This time we performed the NOR, NPR and T-maze tests by applying a retention interval (familiarization→recall period) of 30 min (short-term memory). Strikingly, both sham and SDV mice showed similar recognition performances in all these behavioral paradigms (**Fig. 2A-C** for NOR and **Fig. 2D-F** for NPR), therefore indicating that the vagus nerve is dispensable for short-term recognition memories. Intact short-term spatial working memory and reference memory functions were also observed when sham and SDV mice underwent the Y-maze (**Fig. 2G-I**) and the T-maze tests (**Fig. 2J-K**). Thus, these results indicate that the integrity of the gut-brain vagal axis is not critical for short-term forms of learning and memory.

**Figure 2.**
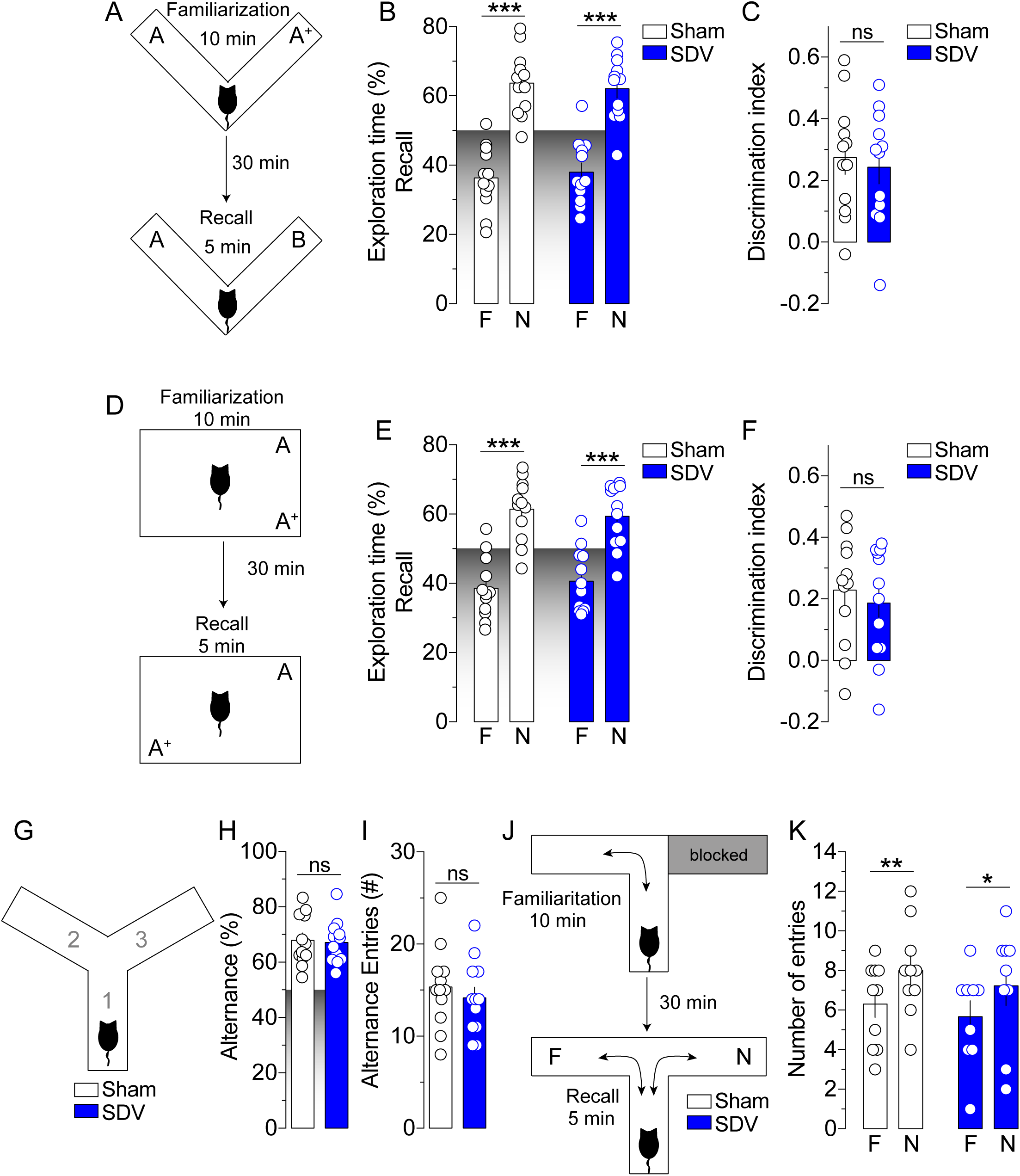
The gut-brain vagal axis is dispensable for short-term recognition memory. (A) The drawing represents the procedure used for the short-term novel object recognition (NOR) test. (B) Percentage of exploration time for sham and SDV mice exploring the familiar (F) and the novel (N) object. (C) Recognition memory discrimination index. (D) The drawing represents the procedure used for the short-term novel place recognition (NPR) test. (E) Percentage of exploration time for sham and SDV mice exploring the familiar (F) and the novel (N) places where objects have been displaced. (F) Recognition memory discrimination index. (G) The drawing indicates the working memory-related Y-Maze test. (H) Percentage of alternance and (I) number of alternance entries of sham and SDV mice during the Y-Maze test. (J) The drawing represents the procedure used to test short-term reference spatial memory by the T-Maze test. (K) Number of entries in the familiar and new compartment. Statistics: *p<0.05, **p<0.01, ***p<0.001 for Sham^(F)^ *vs* Sham^(N)^ or SDV^(F)^ *vs* SDV^(N)^ (B, E, K). For number of mice/group and statistical details see **Suppl. Table 2**.

### The integrity of the gut-brain vagal axis is necessary for hippocampal molecular activity

Several studies have shown that long-term memories also rely on the efficiency of cell signaling events within the hippocampus (Gangarossa et al., 2014a; Kelly et al., 2003; Pettit et al., 2022). Indeed, an alteration of these molecular events may drive to behavioral (mal)adaptations. Thus, here we decided to investigate whether the disruption of the gut-brain vagal axis was followed by a (mal)adapted molecular activity in the hippocampus, notably in the dentate gyrus (DG) which represents the main structural entry of memory-related information. We used two main molecular targets known to be essential for learning and memory processes and whose expression represents a *bona fide* marker of neuronal activity: the immediate early gene (IEG) cFos and the phosphorylated ribosomal protein S6 (Biever et al., 2015; Knight et al., 2012).

We observed that the DG of SDV mice, as compared to sham controls, was characterized by a reduced number of activated p-S6-(**Fig. 3A, B**) and cFos-positive (**Fig. 3D, E**) neurons, indicating that the integrity of the gut-brain vagal tone is necessary for the spontaneous molecular activity of DG-neurons. Interestingly, no major differences in p-S6- and cFos-positive neurons between sham and SDV mice were observed in the hilus (**Fig. 3C, F**), suggesting that gut-brain vagal tone contributes to the spontaneous cellular activity occurring within the DG granule cells layer.

**Figure 3.**
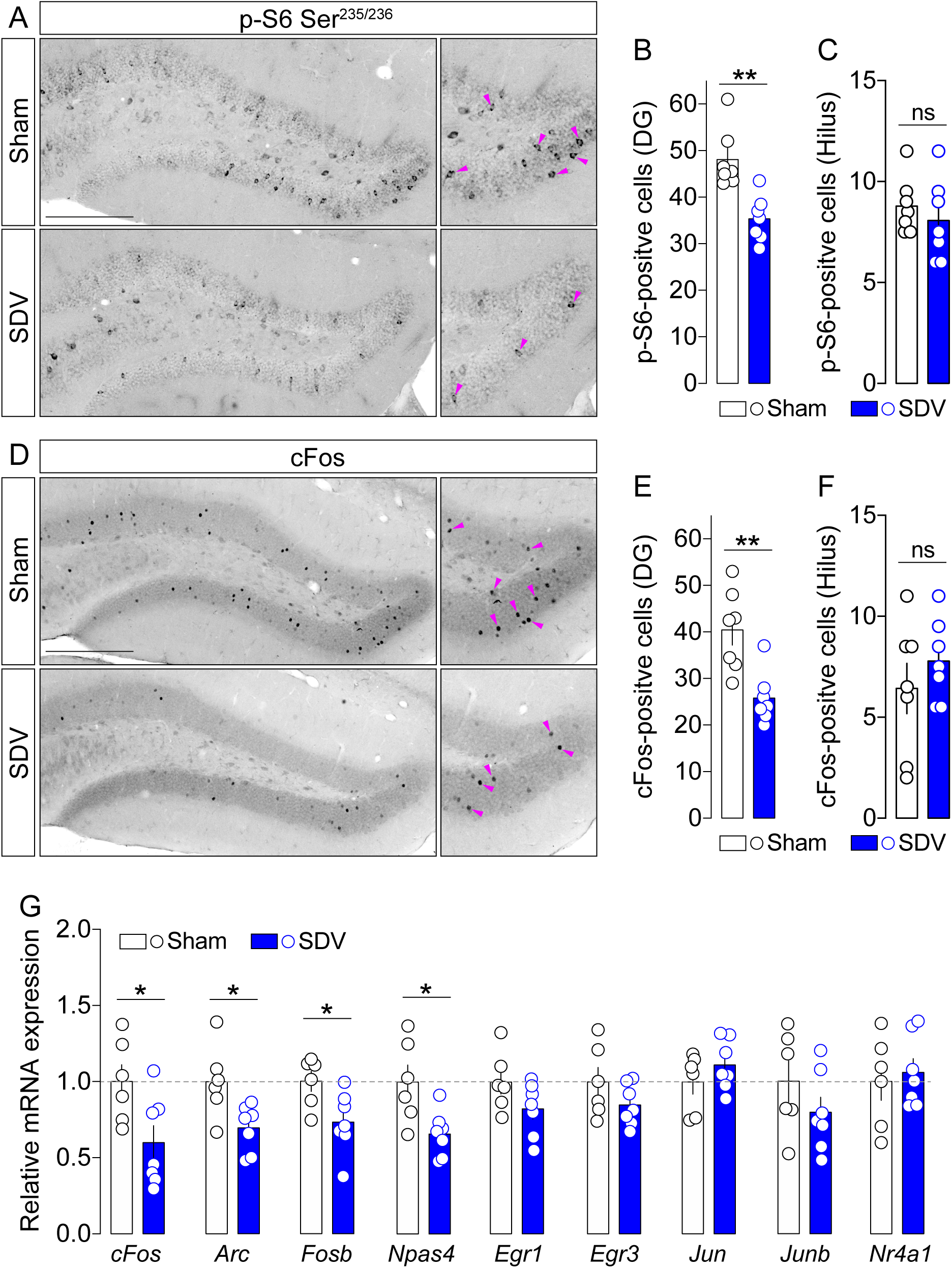
The endogenous gut-brain vagal tone contributes to the molecular activity of the hippocampus. (A) Immunofluorescence detection of rpS6 phosphorylation in the DG of sham and SDV mice (scale bar: 250 μm). Histograms show the number of phospho-rpS6-positive cells in the granular layer (B) and hilus (C) of the DG. (D) Immunofluorescence detection of cFos in the DG of sham and SDV mice (scale bar: 250 μm). Histograms show the number of cFos-positive cells in the granular layer (E) and hilus (F) of the DG. (G) Relative mRNA expression of several immediate early genes such as *cFos*, *Arc*, *Fosb*, *Npas4*, *Egr1*, *Egr3*, *Jun*, *Junb* and *Nr4a1*. Statistics: **p<0.01 for Sham *vs* SDV mice (B and E). Statistics: ***p<0.001 for Sham *vs* SDV (G, for *cFos*, *Arc*, *Fosb* and *Npas4*). For number of mice/group and statistical details see **Suppl. Table 2**.

These results prompted us to investigate whether the disruption of the gut-brain vagal axis led to differential expression of key genes involved in molecular dynamics of plasticity. First, we confirmed the reduction of *cFos* in the hippocampus of SDV mice (**Fig. 3G**). Interestingly, this was also accompanied by a reduction of *Arc*, *Fosb* and *Npas4* (**Fig. 3G**), other key immediate early genes (IEGs) highly involved in learning and memory processes (Chia and Otto, 2013; Gao et al., 2018; Sun and Lin, 2016). However, we also noticed that, despite some trends, other plasticity-related mRNAs (*Egr1*, *Egr3*, *Jun*, *Junb*, *Nr4a1*) were not altered in SDV mice as compared to Sham animals (**Fig. 3G**), indicating that the endogenous vagal tone is required for the basal expression of some but not all IEGs.

These results indicate that the integrity of the gut-brain vagal axis plays a permissive role in modulating the basal molecular activity of the hippocampus and the endogenous expression of key plasticity-associated genes.

### The vagal tone contributes in orchestrating dendritic spines density in the hippocampus

Given our molecular results and the pivotal role of dendritic spines in the integration of synaptic signals and memory traces, we decided to investigate whether the structural morphology and density of hippocampal dendritic spines depended, at least in part, on the integrity of the gut-brain vagal tone. Therefore, we performed 3D diolistic reconstructions of hippocampal dendrites in the dorsal DG and CA1 (**Fig. 4A, F**), the entry and exit regions of the hippocampal circuit, respectively.

**Figure 4.**
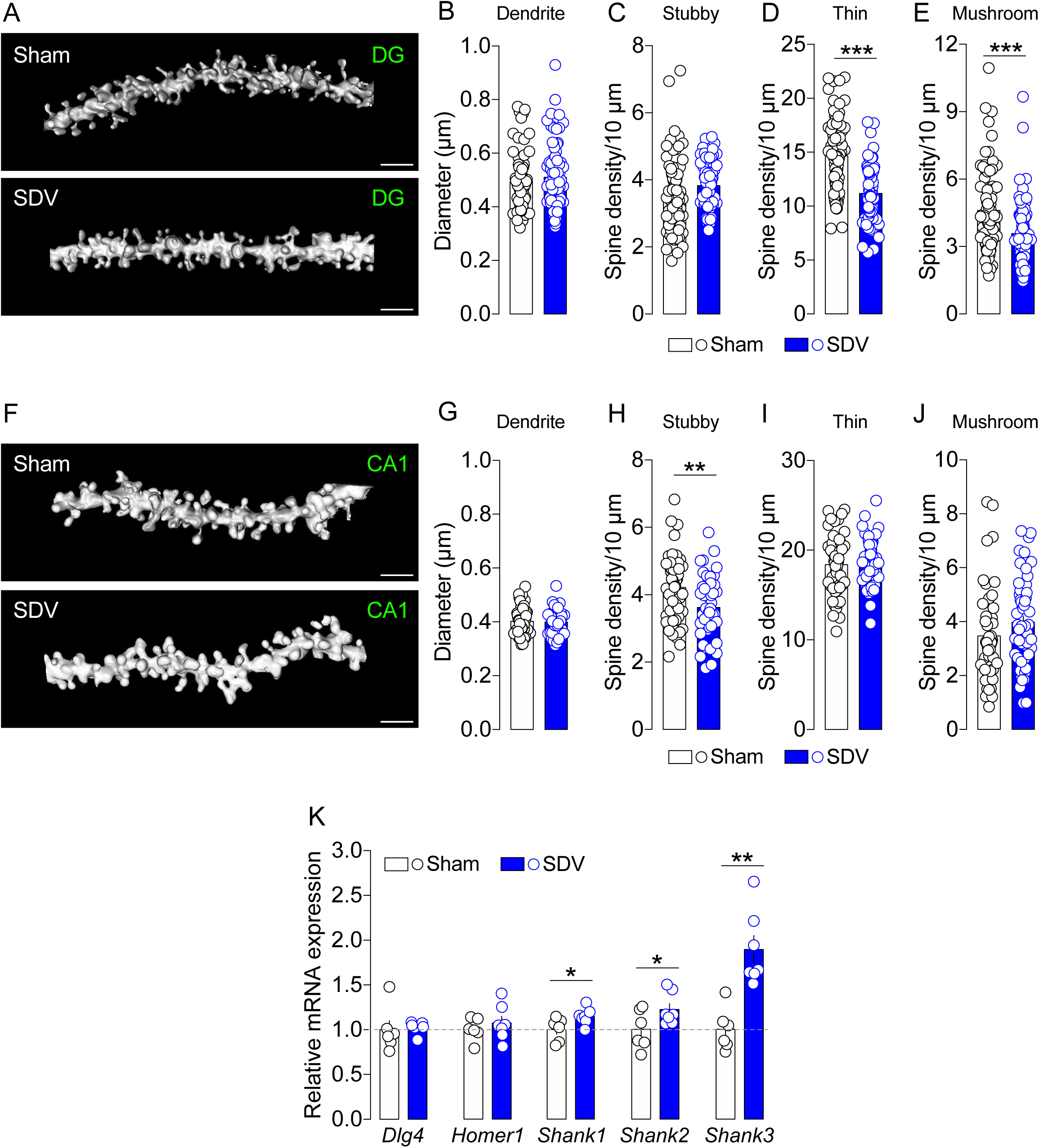
The gut-brain vagal tone shapes the density of hippocampal synaptic spines. (A, F) 3D rendering of dendritic spines in the DG and CA1 (scale bar: 2 μm). Analyses of the dendritic diameter (B, G), density of stubby (C, H), thin (D, I) and mushroom (E, J) spines in the DG and CA1 of sham and SDV mice. (K) Relative mRNA expression of *Dlg4*, *Homer1*, *Shank1*, *Shank2* and *Shank3* in the hippocampus of sham and SDV mice. Statistics: *p<0.05, **p<0.01 and ***p<0.001 for Sham *vs* SDV (D, E, H, K). For number of mice/group and statistical details see **Suppl. Table 2**.

In the DG and CA1, we observed no differences in the diameter of dendrites of sham and SDV mice (**Fig. 4B, G**). Then we analyzed the morphology and density of dendritic spines. Since dendritic spines can be subclassified in different subtypes (*i.e.* stubby, thin, mushroom), we performed a subtypes-specific analysis. In the DG of SDV mice we observed no differences in the density of stubby-like spines as compared to sham mice (**Fig. 4C**), whereas a strong reduction was detected in thin and mushroom spines (**Fig. 4D, E**). In addition to these changes, when analyzing the CA1 (**Fig. 4F**) we also observed a reduction in the density of dendritic spines, however with a different pattern. In fact, in the CA1 of SDV mice we detected a strong reduction of stubby spines as compared to sham mice (**Fig. 4H**), whereas no major alterations were found in the density of thin and mushroom spines (**Fig. 4I, J**). Furthermore, a 3D segmentation of dendritic spines was performed to analyze the distribution of spine head volumes and neck lengths (**Suppl. Fig. 2A**). These complementary structural analyses revealed a reduction in spine head volumes (**Suppl. Fig. 2B, B^1^, D, D^1^**) with no differences in spine neck lengths (**Suppl. Fig. 2C, C^1^, E, E^1^**), indicating that the reduction in dendritic spines observed in SDV mice translates a structural rearrangement of synaptic contacts. These results reveal that the gut-brain vagal tone plays a permissive role in shaping the morphology of dendritic elements (spines) which are key actors in the integration of synaptic signals.

Dendritic spines represent structural elements which contain a plethora of molecular players ensuring the functionality and efficiency of synaptic transmission. Thus, we decided to investigate whether the molecular pattern of some key functional synaptic actors was altered in SDV mice. We observed no differences in the expression of *Dlg4* (PSD-95) and *Homer1a* (**Fig. 4K**). However, we detected an increase in the expression of *Shank1*, *Shank2* and *Shank3* (**Fig. 4K**) in SDV mice, indicating that the integrity of the vagal axis is also important for the homeostatic regulation of molecular synaptic dynamics [*i.e.* stabilization/maturation ensured by scaffolding proteins as Shank(s)] occurring within dendritic spines, an interesting adaptive feature that may counteract the reduction of spines by enriching the molecular repertoire of the remaining spines.

### The gut-brain vagal axis scales hippocampal synaptic plasticity

The above-mentioned behavioral, molecular and morphological adaptations prompted us to study whether the integrity of the gut-brain vagal axis was essential in fine-tuning long-term and short-term forms of hippocampal plasticity at the Schaffer collateral→CA1 synaptic level (**Fig. 5A**), which represents the final and exit step of the intrahippocampal system (DG→CA3→CA1). Thus, we took advantage of *ex vivo* extracellular field potential recordings to measure the synaptic weight of potential vagus-dependent hippocampal (mal)adaptations. First, by using a PP-LFS protocol (Kemp and Bashir, 2001), we assessed whether long-term depression (LTD) was impacted by SDV. We observed a larger LFS-induced LTD in SDV mice as compared to sham animals (**Fig. 5B, B1**). Second, we measured the long-term potentiation (LTP) triggered by high frequency stimulation (HFS) and observed an increase in HFS-induced LTP in SDV mice as compared to sham animals (**Fig. 5C, C1**). Of note, this increase was already transiently evident also during the earliest LTP phase (5 min after HFS, inset of **Fig. 5C**). Moreover, when analyzing the HFS-induced LTP, we observed that, while the LTP induction/consolidation rapidly stabilized in sham mice as expected, the hippocampus of SDV mice was characterized by an unstable form (positive slope) of LTP consolidation (**Suppl. Fig. 3A**). This may depend, at least in part, on the structural and molecular changes observed at the level of dendritic spines (**Fig. 4** and **Suppl. Fig. 2**). Importantly, before and after LTP recordings we performed a paired-pulse facilitation (PPF) protocol to measure short-term forms of plasticity. During baseline (pre-conditioning), we observed a reduction in PPF in SDV mice which, by being inversely related to the probability of quantal glutamate release (Debanne et al., 1996), suggests an impaired short-term potentiation plasticity (**Fig. 5D**). In addition, after the establishment of LTP (post-conditioning) we also observed key differences between the two experimental groups. Notably, while a reduced PPF was measured in sham mice (**Fig. 5D**), an index of presynaptic contribution to LTP (Bliss and Collingridge, 2013; Schulz et al., 1994), no differences in PPF were observed in SDV mice (**Fig. 5D**), suggesting that in this condition global LTP mostly relies on the postsynaptic locus. Of note, in all groups (sham *vs* SDV) and conditions (pre- *vs* post-LTP) facilitation was always observed (PPF>1, **Fig. 5D**), therefore indicating a presynaptic adaptation rather than a distorted synaptic transmission.

**Figure 5.**
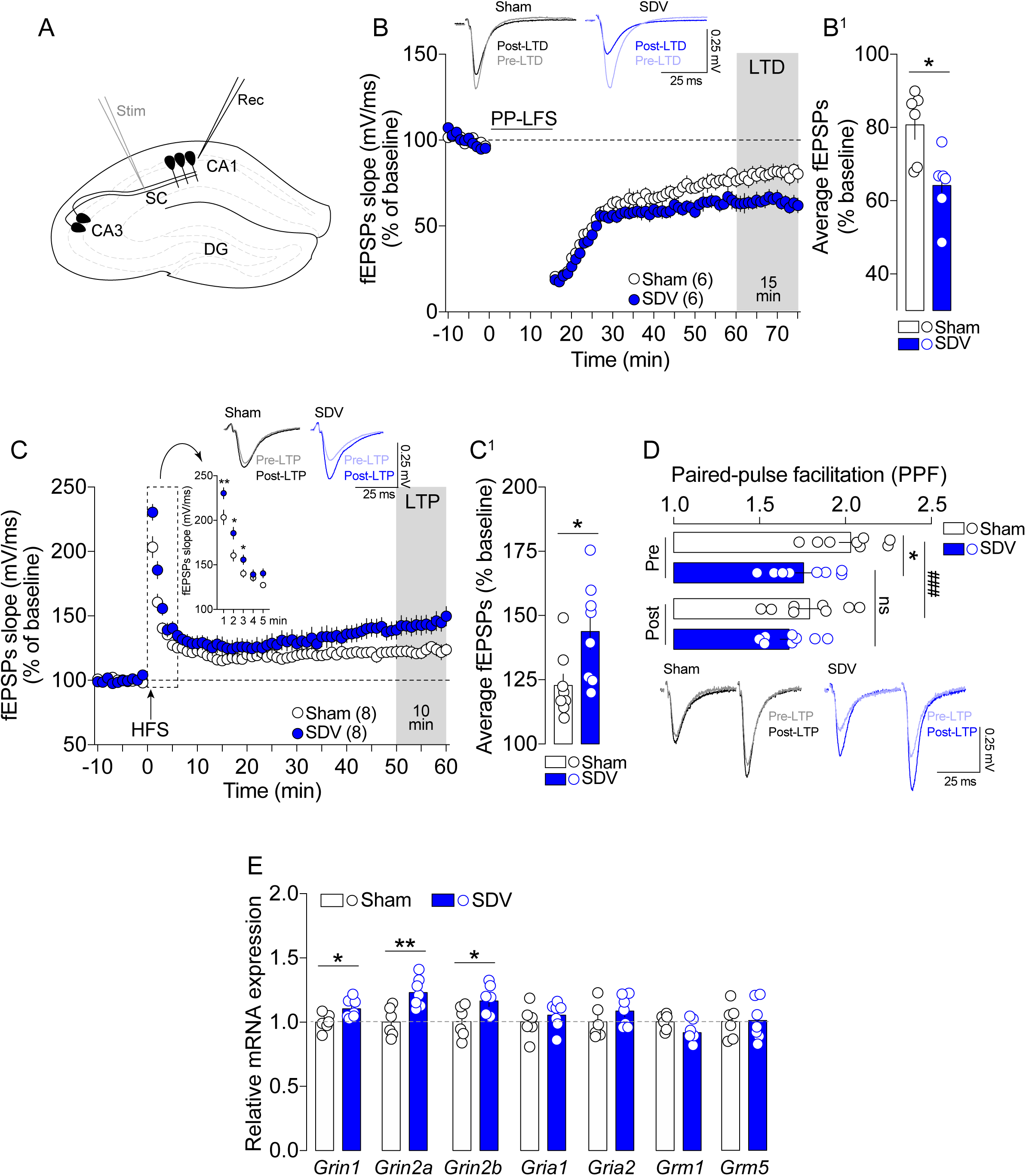
The gut-brain vagal tone governs the dynamics of hippocampal synaptic plasticity. (A) The anatomical drawing shows the *ex vivo* electrophysiological recordings at the level of Schaffer collaterals-to-CA1 synapses. (B) Time-course, representative traces and (B^1^) averaged fEPSPs responses during a PP-LFS-induced long-term depression (LTD) in sham and SDV mice. (C) Time-course, representative traces and (C^1^) averaged fEPSPs responses during an HFS-induced long-term potentiation (LTP) in sham and SDV mice. (D) Paired-pulse facilitation (PPF) and representative traces during baseline (pre) and post-LTP in sham and SDV mice. (E) Relative mRNA expression of glutamate receptors-related subunits such as *Grin1*, *Grin2a*, *Grin2b*, *Gria1*, Gria2, *Grm1* and *Grm5* in the hippocampus of sham and SDV mice. Statistics: *p<0.05 and **p<0.01 for Sham *vs* SDV (B^1^, inset of C, C^1^, D, E). Statistics: ^###^p<0.001 for Sham^(Pre-LTP)^ *vs* Sham^(Post-LTP)^ in the PPF experiment. For number of mice/group and statistical details see **Suppl. Table 2**.

Since hippocampal LTD and LTP strongly rely on glutamate transmission and signaling (Lüscher and Malenka, 2012), we investigated whether SDV was followed by rearrangements in the expression of ionotropic and metabotropic glutamate receptors. Interestingly, in SDV mice we detected a significant increase in different subunits composing the ionotropic NMDA receptor, notably *Grin1* (NR1), *Grin2a* (NR2A) and *Grin2b* (NR2B) (**Fig. 5E**). On the contrary, no alterations were detected for *Gria1* and *Gria2*, two main subunits of the ionotropic AMPA receptor, and for *Grm1* and *Grm5*, two major metabotropic glutamate receptors (mGluRs) (**Fig. 5E**). The increase in *Grin1*, *Grin2a* and *Grin2b* is of interest as NMDA receptor-associated signaling is involved in both LTD and LTP (Lüscher and Malenka, 2012). Since changes in the strength of inhibition (GABA signaling) can directly affect activity-dependent excitatory plasticity (Castillo et al., 2011; Valtcheva et al., 2017; Wang and Maffei, 2014), we analyzed the expression of several GABA-A receptor subunits (α1, α2, α5, β1, β3 and γ2). None of these subunits was altered in SDV mice (**Suppl. Fig. 4A**), thus pointing to glutamate signaling as the main trigger of altered/enlarged LTD and LTP.

These results indicate that the endogenous vagal tone is essential in scaling and shifting short- and long-term forms of hippocampal synaptic plasticity which, by representing cellular forms of learning and memory processes, may altogether contribute to the retention and establishment of memory-like behavioral performances.

## Discussion

Learning and memory functions have largely been attributed to brain structures which, by short- and long-range forms of communication, generate functional and neuroanatomical ensembles. However, the brain is constantly receiving and integrating internal stimuli that, by signaling the physiological states of peripheral organs (interoception), may also influence the elaboration of cognitive functions (*i.e.* learning and memory). This contributes to the establishment of complex and highly interactive body→brain→body systems which regulate different adaptive and/or maladaptive features which cannot be unequivocally attributed to single structures, circuits or neurochemical mediators.

One of the most important, efficient and rapid systems in functionally and bidirectionally connecting the periphery and the brain is represented by the vagus nerve. Our results reveal that the gut-brain vagal system, constitutively and spontaneously, scale hippocampus-dependent learning and memory functions. In fact, we observed that the integrity of the gut-brain vagal axis plays a key and permissive role in scaling behavioral, molecular, cellular and functional ensembles that were essential in shaping learning and memory processes. Notably, we found that disruption of the gut-brain vagal axis impaired long-term, but not short-term, recognition memories while sparing other forms of long-term memory (*e.g.* motor, procedural, reference and working memories). In addition, we observed that those behavioral impairments were accompanied by compelling hippocampal signatures such as a reduced expression of some immediate-early genes (cFos, Arc, Npas4, FosB), an altered morphology of dendritic spines and an unbalanced synaptic plasticity (LTD and LTP). These (mal)adaptive signatures operate as substrates and/or functional proxies to sustain learning and memory (dys)functions.

Seminal articles have already shown that electrical stimulation of the whole vagus nerve (cervical level) improves learning and memory functions in both humans and rodents (Clark et al., 1999, 1998; Olsen et al., 2022; Peña et al., 2014; Sanders et al., 2019) by potentially activating the hippocampus (Xu et al., 2023) and/or by promoting hippocampal neurogenesis and plasticity-associated features (Biggio et al., 2009; Follesa et al., 2007; Olsen et al., 2022; Sanders et al., 2019). In addition, more recent studies have highlighted a clear impact of specific gut-brain vagal manipulations (selective ablations) onto hippocampal functions (O’Leary et al., 2018; Suarez et al., 2018), thus indicating that the gut represents, at least in part, a major peripheral organ with a strong vagus-mediated action onto learning and memory dynamics. To date, the gut-mediated influence onto hippocampal functions has mainly been studied by focusing on the role played by the microbiota whose metabolites, either by acting on vagal afferents or by systemic forms of communication, can contribute in shaping learning and memory (for a recent review see Kuijer and Steenbergen, 2023). However, very little is known regarding the direct role of the gut-brain vagal axis in learning and memory ensembles. Indeed, a few recent and important studies have shown that disruption of the gut-brain vagal axis leads to an alteration of some memory-related behavioral paradigms, a reduction of BDNF and an impairment in the proliferation and survival of newly born hippocampal cells (O’Leary et al., 2018; Suarez et al., 2018). Our results are in line and further expand these findings by revealing that the constitutive gut-brain vagal tone functionally and steadily contributes in shaping hippocampal ensembles. In particular, we show that long-term recognition memory (novel object and novel place recognition), a cognitive function boosted by vagal stimulation even in humans [VNS, (Clark et al., 1999; Giraudier et al., 2020)], strongly relies on the integrity of the gut-brain vagal axis since SDV mice were characterized by a low discriminatory index. Importantly, this impairment was not generalized to other forms of long-term memory and, more interestingly, it did not affect short-term dynamics of recognition memory. This time-dependent difference suggests that the gut-brain vagal tone may not be critical for learning or memory acquisition itself but rather for memory maintenance or storage.

In addition to the behavioral repertoire, our results highlight a key role of the gut-brain vagal axis in orchestrating the molecular and cellular functioning of the hippocampus. First, in SDV mice we observed a reduction of two activity-dependent molecular markers, notably cFos and phospho-ribosomal protein S6 (p-rpS6). Of note, the reduction of p-rpS6, whose activity is usually proportional to mTOR/TOP-dependent mRNA translation (for review see Biever et al., 2015) and highly dependent on a wide range of stimuli (Gangarossa et al., 2014a; Knight et al., 2012; Santini et al., 2012), may underlie a reduced turnover of protein synthesis as well as several molecular/cellular/functional adaptations (Onimus et al., 2022; Puighermanal et al., 2017), therefore contributing in altering key cellular functions.

Strikingly, we also found that the integrity of the gut-brain vagal tone was essential in orchestrating the structural remodeling of hippocampal dendritic spines. Dendritic spines, either mature or immature ones, are neuronal processes which, by undergoing adaptive changes, thoroughly and dynamically contribute in gating synaptic functions and learning and memory ensembles (Berry and Nedivi, 2017; Segal, 2017). Overall, in SDV mice we observed a reduction of dendritic spines in both DG and CA1 hippocampal regions. However, this reduction was heterogeneously distributed within the different types of dendritic spines (*i.e.* mushroom, stubby, thin). In fact, in SDV mice each hippocampal subregion was characterized by a pattern of synaptic modifications with a reduction of mushroom and thin spines in the DG and a reduction of stubby spines in the CA1. While the causative role of spines’ structural remodeling in learning and memory is still debated, a strong positive correlation has been observed between spines’ dynamics (*i.e.* formation, elimination, retraction, stabilization, enlargement) and learning and memory functions (Berry and Nedivi, 2017; Ripoli et al., 2023; Segal, 2017), especially in the hippocampus. Although we cannot provide evidence on whether SDV-induced reduction of mushroom, thin and stubby spines is the result of increased elimination and/or decreased formation/maturation of spines, it is reasonable to hypothesize that a dampened number of dendritic spines following SDV may contribute in shifting and scaling synaptic plasticities (LTD, LTP and PPF). In fact, hippocampal LTD has classically been depicted as leading to a shrinkage, retraction or loss of dendritic spines (Bastrikova et al., 2008; Hasegawa et al., 2015; Zhou et al., 2004). Therefore, it would be reasonable to expect that a basal reduction of dendritic spines or a slower maturation of thin/stubby spines into mushroom spines may lead to an enhancement of LTD, as observed in SDV mice.

Contrary to LTD, hippocampal LTP is classically associated to increased density, maturation and stabilization of dendritic spines (Runge et al., 2020). However, SDV mice showed a facilitated LTP despite a lower number of spines. While at first these cellular features may sound counterintuitive, it is worth to mention that SDV mice showed an unstable LTP induction/consolidation (positive slope) as well as an altered PPF at the basal level followed by a lack of PPF adaptation after LTP induction, thus showing reduced adaptability at least at the presynaptic locus. Moreover, this facilitated form of LTP may be associated to a pre-existing SDV-induced (mal)adaptive adjustments and enrichment of Shank1, Shank2 and Shank3, well-known scaffolding actors of mature spines involved in synaptic transmission and LTP (Jaramillo et al., 2016; Kouser et al., 2013). In fact, we observed that disruption of the gut-brain vagal axis led to a significant increase of Shank(s), which may therefore further facilitate the establishment of LTP. This is further highlighted by the fact that Shank3 functionally contributes to the glutamate receptosome by participating to the regulation of NMDA receptors’ activity (Duffney et al., 2013; Kouser et al., 2013). In fact, we observed that the expression of key plasticity-associated NMDA subunits (Grin1, Grin2a and Grin2b) was increased in SDV mice, therefore indicating a rearrangement of glutamate transmission which may enlarge the window of NMDA receptor-dependent LTP and LTD (Lüscher and Malenka, 2012). It may be reasonable to speculate that a reduction in dendritic spines, as observed when the gut-brain vagal integrity is compromised, may be accompanied by a dynamic molecular process that, by enhancing Shank(s)/NMDAR functions, tries to optimize the efficiency of synaptic contacts which, despite this rearrangement, cannot sustain long-term engrams or memory traces.

It is important to mention that, although an increased LTP is usually thought to mimic more efficient learning and memory functions, other studies have shown detrimental memory and learning functions despite enhanced LTP (Brun et al., 2001; Jolas et al., 2002; Kim et al., 2009; Migaud et al., 1998; Navakkode et al., 2022; Vaillend et al., 2004). Hence, we argue that the larger shift towards opposite forms of plasticity (enhanced LTD and LTP) coupled to the concomitant impaired short-term facilitation as well as to the aberrant rearrangements of dendritic spines may functionally and collectively lead to selective learning and memory impairments (recognition memory *vs* other forms of memory) in SDV mice.

The impairments in long-term recognition memory observed in SDV mice are also supported by a reduction in the basal expression of immediate early genes (IEGs) such as *cFos*, *Arc*, *Npas4* and *Fosb*. In fact, it is well-known that the expression and/or induction of IEGs is a critical process underlying long-term learning and memory functions (Minatohara et al., 2015). Indeed, a reduced pool of readily available *cFos*, *Arc*, *Npas4* and *Fosb* may lead to a reduced storage of long-term memory traces or engrams which may in turn drive to behavioral memory deficits.

Importantly, our results should be put in relationship with emerging studies suggesting that even neurodegenerative disorders (Parkinson’s and Alzheimer’s diseases), which are characterized by profound learning and memory impairments, seem to be mediated, at least in part, by a dysfunctional gut-brain vagal axis. In fact, the vagal axis contributes to the neuronal seeding and spreading of neurodegenerative hallmarks (misfolded α-synuclein and β-amyloid proteins) from the gut to the brain (Braak et al., 2003; Chandra et al., 2023; Chen et al., 2021; Kim et al., 2019; Sun et al., 2020), therefore potentially participating to the cognitive decline observed in neurodegenerative disorders. In addition, other breakthroughs have shown that the vagus nerve, as a functional bridge between the gut microbiota and the brain, can mediate some forms of dysbiosis-induced cognitive and neurological dysfunctions (Kuijer and Steenbergen, 2023; Spichak et al., 2021).

In conclusion, our results reveal that the gut-brain vagal axis exerts a strong long-range interoceptive control over different hippocampal dynamics which may converge onto altered cognitive performances. This compelling evidence contributes in expanding the conceptual framework according to which cognitive functions tightly rely on a correct and well-balanced body-brain communication and may pave the way to new forms of therapeutic approaches aimed at restoring learning and memory dysfunctions by leveraging the power of the gut-brain axis.

## Acknowledgments

We thank Olja Kacanski for administrative support, Isabelle Le Parco, Daniel Quintas, Magguy Boa, Ludovic Maingault and Angélique Dauvin for animals’ care. We thank Charles Le Ciclé for technical advises. We acknowledge the Functional and Physiological Exploration platform (FPE) of the Université Paris Cité (UPCité), CNRS, Unité de Biologie Fonctionnelle et Adaptative, and the animal core facility “Buffon” of UPCité/Institut Jacques Monod.

## Funding

This work was supported by the Nutricia Research Foundation (#2022-E7), Agence Nationale de la Recherche (ANR-21-CE14-0021-01, ANR-23-CE14-0014-02), Fédération pour la Recherche sur le Cerveau and Association France Parkinson, Institut universtaire de France, Université Paris Cité (Stratex_Idex-2023-028 EMERGENCE), and CNRS. O.O. is supported by an FRM Ph.D fellowship. I.N.O.S and J-P. M. were supported by an EMBO long-term postdoctoral fellowship (ALTF 1981-2020 to I.N.O.S), ENS Paris-Saclay, CNRS and Université Paris-Saclay.

## Competing interests

The authors declare no competing interests.

## CRediT authorship contribution statement

**Oriane Onimus**: Conceptualization, Methodology, Validation, Formal analysis, Investigation, Data Curation, Writing - Review & Editing, Visualization. **Faustine Arrivet**: Methodology, Formal analysis, Investigation, Data Curation. **Isis Nem de Oliveira Souza**: Methodology, Formal analysis, Investigation, Data Curation, Writing - Review & Editing. **Benoit Bertrand**: Formal analysis, Investigation. **Julien Castel**: Methodology. **Serge Luquet**: Writing - Review & Editing. **Jean-Pierre Mothet**: Formal analysis, Data Curation, Supervision, Writing - Review & Editing. **Nicolas Heck**: Methodology, Formal analysis, Investigation, Data Curation, Supervision, Writing - Review & Editing. **Giuseppe Gangarossa**: Conceptualization, Formal analysis, Investigation, Data Curation, Resources, Writing - Original Draft, Writing - Review & Editing, Project administration, Funding acquisition.

## Figure legends

**Suppl. Figure 1. The gut-brain vagal axis is not critical for other forms of long-term memory.** (A) Latency to fall (sec) during the rotarod test over 4 training days (D1 to D4) in sham and SDV mice. Inset indicates learning and memory motor performances during Day1 (D1). (B, B^1^) Locomotor activity, kinetics for B and cumulative activity for B^1^, of sham and SDV mice exposed to a novel environment over 2 consecutive days (bb for beam breaks). Statistics: ***p<0.001 for Sham^(trial3_D1)^ *vs* Sham^(trial1_D1)^ or SDV^(trial3_D1)^ *vs* SDV^(trial1_D1)^ (inset of A). Statistics: ***p<0.001 for Sham^(D2)^ *vs* Sham^(D1)^ or SDV^(D2)^ *vs* SDV^(D1)^ (B^1^). For number of mice/group and statistical details see **Suppl. Table 2**.

**Suppl. Figure 2. The gut-brain vagal axis is not critical for spine head volumes and spine lengths.** (A) 3D rendering of a dendritic segment of the DG and the segmented spine heads (scale bar: 2 μm). Segmentation procedure allows the measurement of head volumes in 3D from spines. (B-B^1^, D-D^1^) Frequency distribution and averages of spine head volumes in the DG (B, B^1^) and CA1 (D, D^1^). (C-C^1^, E-E^1^) Frequency distribution and averages of spine neck lengths, calculated as the distance between the center of the head and the border of the dendrite, in the DG (C, C^1^) and CA1 (E, E^1^). Statistics: ***p<0.001 and **p<0.01 for SDV *vs* Sham. For statistical details see **Suppl. Table 2**.

**Suppl. Figure 3. Disruption of the gut-brain vagal axis alters LTP consolidation.** (A) Comparisons of fEPSPs responses during LTP induction/consolidation at two intervals (30-40 min and 50-60 min post-HFS). Statistics: **p<0.01 for SDV^50-60min^ *vs* SDV^30-40min^. For number of mice/group and statistical details see **Suppl. Table 2**.

**Suppl. Figure 4. The gut-brain vagal axis is not critical for the expression of GABA-A receptor’s subunits.** (A) Relative mRNA expression of GABA-A α1, α2, α5, β1, β3 and γ2 subunits in the hippocampus of sham and SDV mice. For number of mice/group and statistical details see **Suppl. Table 2**.

## Notes

### Competing Interest Statement

The authors have declared no competing interest.

