## Supplementary figures and images for "The gut-brain vagal axis scales hippocampal memory processes and plasticity"

### Suppl. Figure 1

#### Suppl. Figure 1

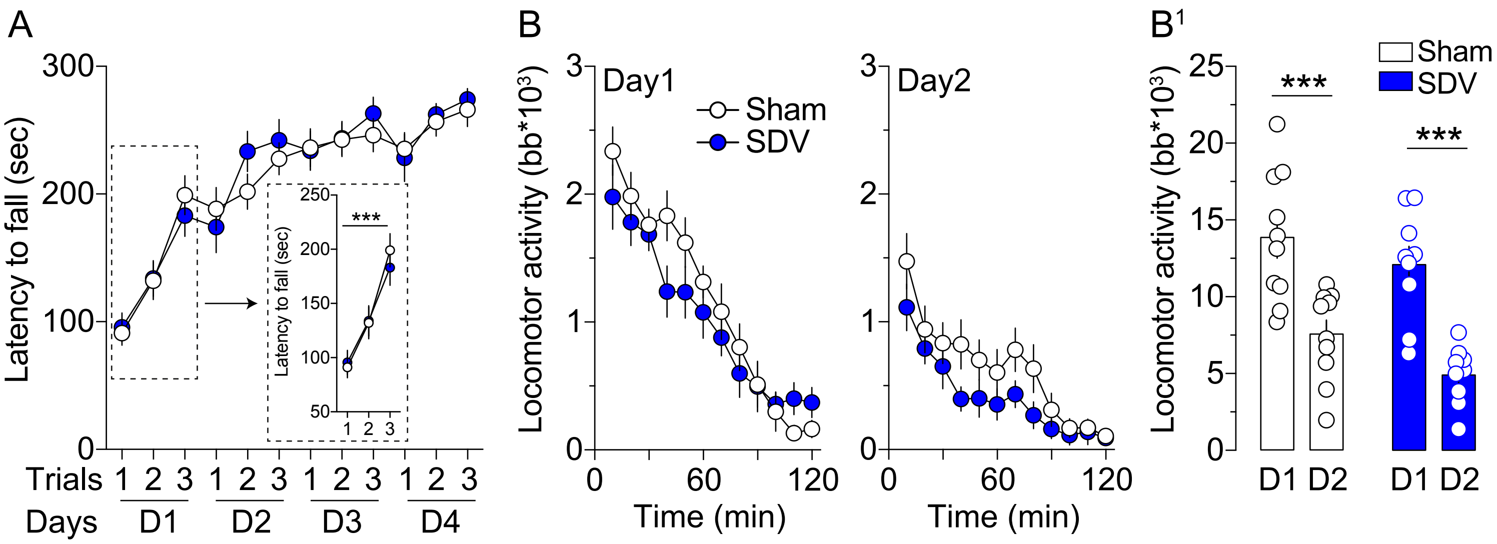

### Suppl. Figure 2

#### Suppl. Figure 2

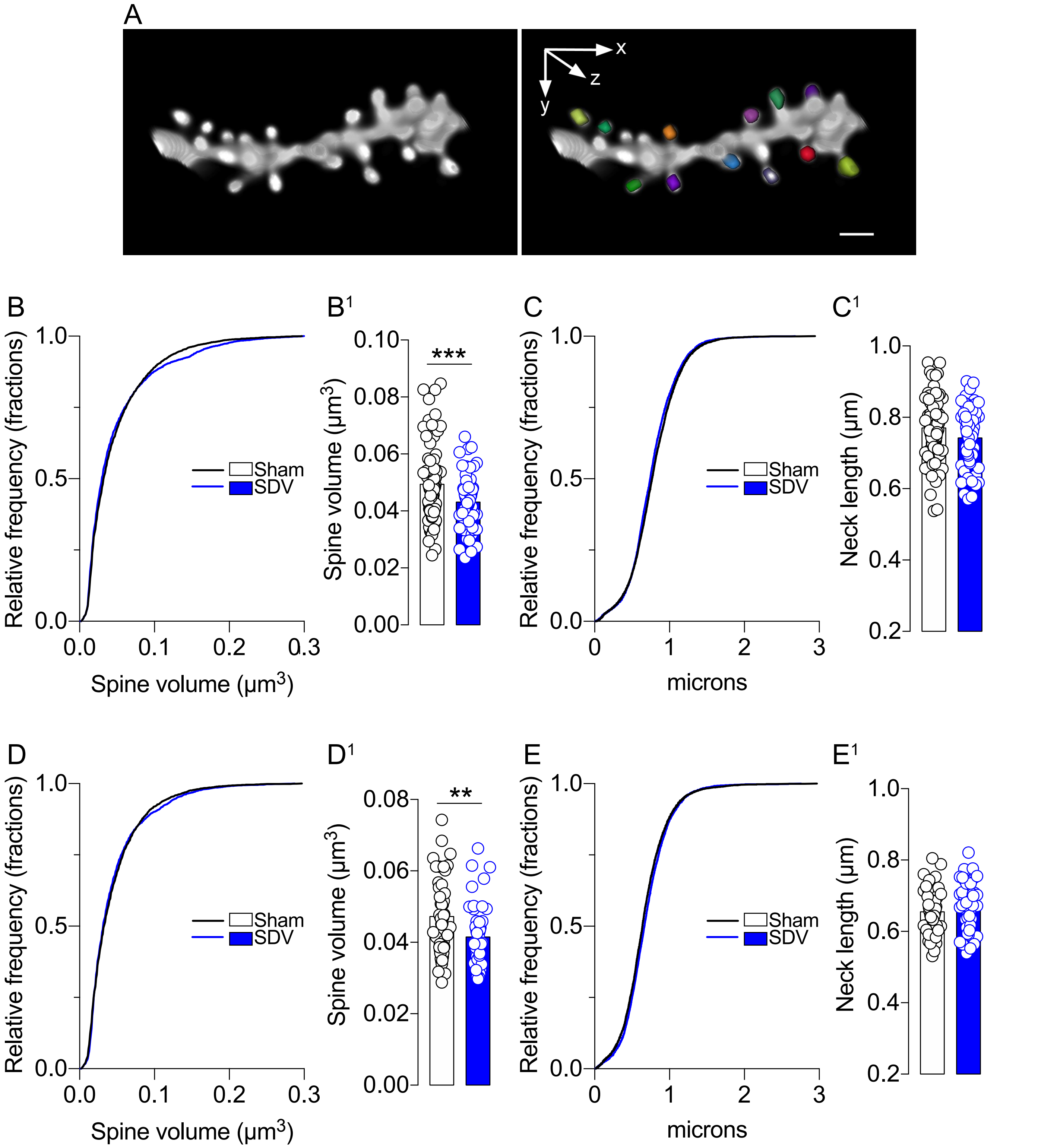

### Suppl. Figure 3

#### Suppl. Figure 3

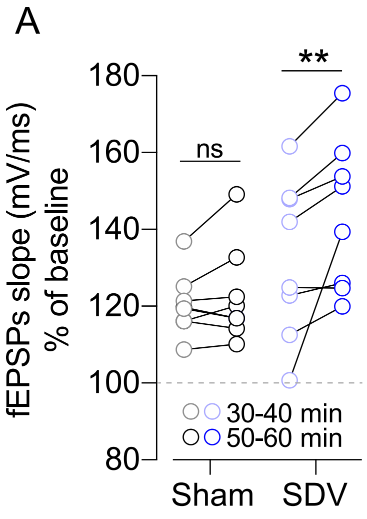

### Suppl. Figure 4

#### Suppl. Figure 4

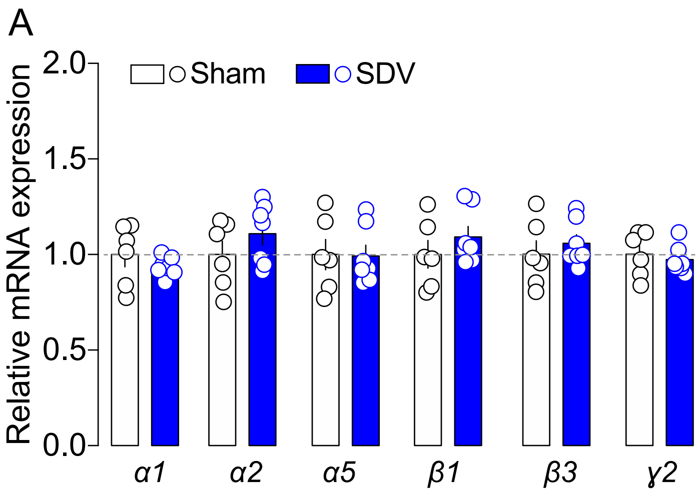
