## Supplementary material for "The gut-brain vagal axis scales hippocampal memory processes and plasticity": Suppl. Table 1

| **Primers of Figs. 3, 4, 5 and Suppl. Fig. 4** | | |
|---|---|---|
| **Gene** | **Sens** | **5' → 3'** |
| Arc | Forward | AAGTGCCGAGCTGAGATGC |
| | Reverse | CGACCTGTGCAACCCTTTC |
| Dlg4 | Forward | TGAGATCAGTCATAGCAGCTACT |
| | Reverse | CTTCCTCCCCTAGCAGGTCC |
| Egr1 | Forward | CGAACAACCCTATGAGCACCTG |
| | Reverse | CAGAGGAAGACGATGAAGCAGC |
| Egr3 | Forward | CCGGTGACCATGAGCAGTTT |
| | Reverse | TAATGGGCTACCGAGTCGCT |
| Fos | Forward | CGGGTTTCAACGCCGACTA |
| | Reverse | TTGGCACTAGAGACGGACAGA |
| Fosb | Forward | TTTTCCCGGAGACTACGACTC |
| | Reverse | GTGATTGCGGTGACCGTTG |
| α1(Gabra1) | Forward | AAAAGCGTGGTTCCAGAAAA |
| | Reverse | GCTGGTTGCTGTAGGAGCAT |
| α2 (Gabra2) | Forward | GCTACGCTTACACAACCTCAGA |
| | Reverse | GACTGGCCCAGCAAATCATACT |
| α5 (Gabra5) | Forward | GATTGTGTTCCCCATCTTGTTTGGC |
| | Reverse | TTACTTTGGAGAGGTGGCCCCTTTT |
| β1 (Gabrb1) | Forward | GGTTTGTTGTGCACACAGCTCC |
| | Reverse | ATGCTGGCGACATCGATCCGC |
| β3 (Gabrb3) | Forward | GAGCGTAAACGACCCCGGGAA |
| | Reverse | GGGACCCCCGAAGTCGGGTCT |
| γ2 (Gabrg2) | Forward | ACTTCTGGTGACTATGTGGTGAT |
| | Reverse | GGCAGGAACAGCATCCTTATTG |
| Gria1 | Forward | GTCCGCCCTGAGAAATCCAG |
| | Reverse | CTCGCCCTTGTCGTACCAC |
| Gria2 | Forward | GCCGAGGCGAAACGAATGA |
| | Reverse | CACTCTCGATGCCATATACGTTG |
| Grin1 | Forward | AGAGCCCGACCCTAAAAAGAA |
| | Reverse | CCCTCCTCCCTCTCAATAGC |
| Grin2a | Forward | ACGTGACAGAACGCGAACTT |
| | Reverse | TCAGTGCGGTTCATCAATAACG |
| Grin2b | Forward | GCCATGAACGAGACTGACCC |
| | Reverse | GCTTCCTGGTCCGTGTCATC |
| Grm1 | Forward | CATACGGAAAGGGGAAGTGA |
| | Reverse | AAAAGGCGATGGCTATGATG |
| Grm5 | Forward | ACGAAGACCAACCGTATTGC |
| | Reverse | AGACTTCTCGGATGCTTGGA |
| Homer1 | Forward | CCCTCTCTCATGCTAGTTCAGC |
| | Reverse | GCACAGCGTTTGCTTGACT |
| Jun | Forward | CCCTCTCTCATGCTAGTTCAGC |
| | Reverse | GCACAGCGTTTGCTTGACT |
| Junb | Forward | TTTTGTCAAAGCCCTGGACG |
| | Reverse | GGGGAGTAACTGCTGAGGTT |
| Npas4 | Forward | CTGCATCTACACTCGCAAGG |
| | Reverse | GCCACAATGTCTTCAAGCTCT |
| Nr4a1 | Forward | GTTATCCGAAAGTGGGCAGA |
| | Reverse | AGTACCAGGCCTGAGCAGAA |
| Rpl19 | Forward | GGGCAGGCATATGGGCATA |
| | Reverse | GGCGGTCAATCTTCTTGGATT |
| Shank1 | Forward | CCGCTACAAGACCCGAGTCTA |
| | Reverse | CCTGAATCTGAGTCGTGGTAGTT |
| Shank2 | Forward | AGAGGCCCCAGCTTATTCCAA |
| | Reverse | CAGGGGTATAGCTTCCAAGGC |
| Shank3 | Forward | ATGGGCCTGTGTGGTAGTCTT |
| | Reverse | CCACCTTATCTGTGCTGTGTAG |
