## Supplementary material for "The gut-brain vagal axis scales hippocampal memory processes and plasticity": Suppl. Table 2

### Supplementary Table 2

| Statistics of Figure 1 | | | | | | |
|---|---|---|---|---|---|---|
| Figure panels | | n | Statistical analysis | | F-value | p-value |
| Fig 1B | NOR (Recall) | Sham n=12, SDV n=12 | 2-way ANOVA | interaction | $F_{(1, 44)} = 55.17$ | $p < 0.0001$ |
| | | | | time | $F_{(1, 44)} = 85.14$ | $p < 0.0001$ |
| | | | | groups | $F_{(1, 44)} = 0$ | $p > 0.9999$ |
| Fig 1C | NOR (Recall-Discrimination) | Sham n=12, SDV n=12 | Unpaired t-test | | | $p < 0.0001$ t=5.297, df=22 |
| Fig 1D | NOR (Familiarization) | Sham n=12, SDV n=12 | 2-way ANOVA | interaction | $F_{(1, 44)} = 0.00147$ | $p = 0.9696$ |
| | | | | time | $F_{(1, 44)} = 0.03667$ | $p = 0.8490$ |
| | | | | groups | $F_{(1, 44)} = 1.758$ | $p = 0.1918$ |
| Fig 1F | NPR (Recall) | Sham n=12, SDV n=12 | 2-way ANOVA | interaction | $F_{(1, 44)} = 49.63$ | $p < 0.0001$ |
| | | | | time | $F_{(1, 44)} = 94.57$ | $p < 0.0001$ |
| | | | | groups | $F_{(1, 44)} = 0$ | $p > 0.9999$ |
| Fig 1G | NPR (Recall-Discrimination) | Sham n=12, SDV n=12 | Unpaired t-test | | | $p < 0.0001$ t=5.001, df=22 |
| Fig 1H | NPR (Familiarization) | Sham n=12, SDV n=12 | 2-way ANOVA | interaction | $F_{(1, 44)} = 0.00597$ | $p = 0.9388$ |
| | | | | time | $F_{(1, 44)} = 0.01558$ | $p = 0.9012$ |
| | | | | groups | $F_{(1, 44)} = 0.00199$ | $p = 0.9646$ |
| Fig 1J | T-Maze | Sham n=10, SDV n=9 | 2-way ANOVA | interaction | $F_{(1, 17)} = 0.00017$ | $p = 0.9897$ |
| | | | | time | $F_{(1, 17)} = 20$ | $p = 0.0003$ |
| | | | | groups | $F_{(1, 17)} = 0.7212$ | $p = 0.4076$ |

| Statistics of Figure 2 | | | | | | |
|---|---|---|---|---|---|---|
| Figure panels | | n | Statistical analysis | | F-value | p-value |
| Fig 2B | NOR (Recall) | Sham n=12, SDV n=12 | 2-way ANOVA | interaction | $F_{(1, 44)} = 0.3801$ | $p = 0.5407$ |
| | | | | time | $F_{(1, 44)} = 92.32$ | $p < 0.0001$ |
| | | | | groups | $F_{(1, 44)} = 0$ | $p > 0.9999$ |
| Fig 2B | NOR (Recall-Discrimination) | Sham n=12, SDV n=12 | Unpaired t-test | | | $p = 0.6882$ t=0.4067, df=22 |
| Fig 2E | NPR (Recall) | Sham n=12, SDV n=12 | 2-way ANOVA | interaction | $F_{(1, 44)} = 0.641$ | $p = 0.4277$ |

| | | | | | | |
|---|---|---|---|---|---|---|
| | | | | time | $F_{(1, 44)} = 64.35$ | p < 0.0001 |
| | | | | groups | $F_{(1, 44)} = 0$ | p > 0.9999 |
| Fig 2F | NPR (Recall-Discrimination) | Sham n=12, SDV n=12 | Unpaired t-test | | | p = 0.5755 t=0.5684, df=22 |
| Fig 2H | % of alternance (Y-Maze) | Sham n=12, SDV n=12 | Unpaired t-test | | | p = 0.8245 t=0.2244, df=22 |
| Fig 2I | Alternance entries (Y-Maze) | Sham n=12, SDV n=12 | Unpaired t-test | | | p = 0.5056 t=0.6767, df=22 |
| Fig 2K | T-Maze | Sham n=10, SDV n=9 | 2-way ANOVA | interaction | $F_{(1, 17)} = 0.03627$ | p = 0.8512 |
| | | | | time | $F_{(1, 17)} = 18.42$ | p = 0.0005 |
| | | | | groups | $F_{(1, 17)} = 0.4442$ | p = 0.5141 |

| Statistics of Figure 3 | | | | | |
|---|---|---|---|---|---|
| Figure panels | | n | Statistical analysis | F-value | p-value |
| Fig 3B | pS6-cell in DG | Sham n=7, SDV n=7 | Unpaired t-test | | p = 0.0013 t=4.172, df=12 |
| Fig 3C | pS6-cell in Hilus | Sham n=7, SDV n=7 | Unpaired t-test | | p = 0.4592 t=0.7647, df=12 |
| Fig 3E | cFos-cell in DG | Sham n=7, SDV n=7 | Unpaired t-test | | p = 0.0027 t=3.768, df=12 |
| Fig 3F | cFos-cell in Hilus | Sham n=7, SDV n=7 | Unpaired t-test | | p = 0.3712 t=0.929, df=12 |
| Fig 3G | *cFos* | Sham n=6, SDV n=7 | Unpaired t-test | | p = 0.0278 t=2.533, df=11 |
| | *Arc* | | | | p = 0.0179 t=2.782, df=11 |
| | *Fosb* | | | | p = 0.0231 t=2.638, df=11 |
| | *Npas4* | | | | p = 0.0147 t=2.891, df=11 |
| | *Egr1* | | | | p = 0.1094 t=1.742, df=11 |
| | *Egr3* | | | | p = 0.1777 t=1.44, df=11 |

| | | | | | |
|---|---|---|---|---|---|
| | *Jun* | | | | p = 0.2822<br>t=1.131, df=11 |
| | *Junb* | | | | p = 0.2402<br>t=1.242, df=11 |
| | *Nr4a1* | | | | p = 0.6863<br>t=0.4148, df=11 |

| Statistics of Figure 4 | | | | | |
|---|---|---|---|---|---|
| Figure panels | | n | Statistical analysis | F-value | p-value |
| Fig 4B | Dendritic diameter (DG) | Sham n=80 dendrites from 7 mice,<br><br>SDV n=71 dendrites from 5 mice | Unpaired t-test | | p = 0.2451<br>t=1.167, df=149 |
| Fig 4C | Stubby spines (DG) | Sham n=80 dendrites from 7 mice,<br><br>SDV n=71 dendrites from 5 mice | Unpaired t-test | | p = 0.1229<br>t=1.552, df=149 |
| Fig 4D | Thin spines (DG) | Sham n=80 dendrites from 7 mice,<br><br>SDV n=71 dendrites from 5 mice | Unpaired t-test | | p < 0.0001<br>t=7.002, df=149 |
| Fig 4E | Mushroom spines (DG) | Sham n=80 dendrites from 7 mice,<br><br>SDV n=71 dendrites from 5 mice | Unpaired t-test | | p = 0.0002<br>t=3.761, df=149 |
| Fig 4G | Dendritic diameter (CA1) | Sham n=49 dendrites from 5 mice,<br><br>SDV n=51 dendrites from 5 mice | Unpaired t-test | | p = 0.8453<br>t=0.1956, df=98 |
| Fig 4H | Stubby spines (CA1) | Sham n=49 dendrites from 5 mice,<br><br>SDV n=51 dendrites from 5 mice | Unpaired t-test | | p = 0.0012<br>t=3.342, df=98 |
| Fig 4I | Thin spines (CA1) | Sham n=49 dendrites from 5 mice,<br><br>SDV n=51 dendrites from 5 mice | Unpaired t-test | | p = 0.6562<br>t=0.4465, df=98 |
| Fig 4J | Mushroom spines (CA1) | Sham n=49 dendrites from 5 mice, | Unpaired t-test | | p = 0.1382<br>t=1.495, df=98 |

| | | | | | |
|---|---|---|---|---|---|
| | | SDV n=51 dendrites from 5 mice | | | |
| Fig 4K | *Dlg4* | Sham n=6, SDV n=7 | Unpaired t-test | | p = 0.7960 t=0.2648, df=11 |
| | *Homer1* | | | | p = 0.4586 t=0.7681, df=11 |
| | *Shank1* | | | | p = 0.0413 t=2.31, df=11 |
| | *Shank2* | | | | p = 0.0488 t=2.214, df=11 |
| | *Shank3* | | | | p = 0.0007 t=4.66, df=11 |

**Statistics of Figure 5**

| Figure panels | | n | Statistical analysis | | F-value | p-value |
|---|---|---|---|---|---|---|
| Fig 5B[1] | LTD | Sham n=6, SDV n=6 | Unpaired t-test | | | p = 0.0121 t=3.0591, df=10 |
| Fig 5C inset | LTP | Sham n=8, SDV n=8 | 2-way ANOVA | interaction | $F_{(4, 56)} = 5.941$ | p = 0.0005 |
| | | | | time | $F_{(4, 56)} = 314$ | p < 0.0001 |
| | | | | groups | $F_{(1, 14)} = 5.631$ | p = 0.0325 |
| Fig 5C[1] | LTP | Sham n=8, SDV n=8 | Unpaired t-test | | | p = 0.0230 t=2.5534, df=14 |
| Fig 5D | PPF | Sham n=8, SDV n=8 | 2-way ANOVA | interaction | $F_{(1, 14)} = 4.907$ | p = 0.0438 |
| | | | | time | $F_{(1, 14)} = 22.03$ | p = 0.0003 |
| | | | | groups | $F_{(1, 14)} = 5.397$ | p = 0.0358 |
| Fig 5E | *Grin1* | Sham n=6, SDV n=7 | Unpaired t-test | | | p = 0.0213 t=2.684, df=11 |
| | *Grin2a* | | | | | p = 0.0044 t=3.569, df=11 |
| | *Grin2b* | | | | | p = 0.0337 t=2.426, df=11 |
| | *Gria1* | | | | | p = 0.4292 t=0.8208, df=11 |
| | *Gria2* | | | | | p = 0.2561 t=1.198, df=11 |
| | *Grm1* | | | | | p = 0.0825 t=1.91, df=11 |
| | *Grm5* | | | | | p = 0.9154 t=0.1087, df=11 |

| Statistics of Suppl. Figure 1 | | | | | | |
|---|---|---|---|---|---|---|
| Figure panels | | n | Statistical analysis | | F-value | p-value |
| SFig 1A | Rotarod (from D1 to D4) | Sham n=12, SDV n=12 | 2-way ANOVA | interaction | $F_{(11, 242)} = 0.683$ | p = 0.7541 |
| | | | | time | $F_{(11, 242)} = 43.81$ | p < 0.0001 |
| | | | | groups | $F_{(1, 22)} = 0.08478$ | p = 0.7737 |
| | Rotarod (D1, inset) | | 2-way ANOVA | interaction | $F_{(2, 44)} = 0.5163$ | p = 0.6003 |
| | | | | time | $F_{(2, 44)} = 40.13$ | p < 0.0001 |
| | | | | groups | $F_{(1, 22)} = 0.04786$ | p = 0.8289 |
| SFig 1B | Locomotor activity (Day1) | Sham n=10, SDV n=9 | 2-way ANOVA | interaction | $F_{(11, 187)} = 1.638$ | p = 0.0910 |
| | | | | time | $F_{(11, 187)} = 47.22$ | p < 0.0001 |
| | | | | groups | $F_{(1, 17)} = 0.9232$ | p = 0.3501 |
| | Locomotor activity (Day2) | | 2-way ANOVA | interaction | $F_{(11, 187)} = 0.5992$ | p = 0.8280 |
| | | | | time | $F_{(11, 187)}) = 14.3$ | p < 0.0001 |
| | | | | groups | $F_{(1, 17)} = 5.224$ | p = 0.0354 |
| SFig 1B[1] | Locomotor activity (Day1 *vs* Day2) | Sham n=10, SDV n=9 | 2-way ANOVA | interaction | $F_{(1, 17)} = 0,3792$ | p = 0.5462 |
| | | | | time | $F_{(1, 17)} = 87.84$ | p < 0.0001 |
| | | | | groups | $F_{(1, 17)} = 2.685$ | p = 0.1197 |

| Statistics of Suppl. Figure 2 | | | | | |
|---|---|---|---|---|---|
| Figure panels | | n | Statistical analysis | F-value | p-value |
| SFig 2B[1] | Spine volume in DG | Sham n=78, SDV n=73 | Unpaired t-test | | p = 0.0010 t=3.37, df=149 |
| SFig 2C[1] | Neck length in DG | Sham n=79, SDV n=73 | Unpaired t-test | | p = 0.05 t=2.039, df=150 |
| SFig 2D[1] | Spine volume in CA1 | Sham n=49, SDV n=52 | Unpaired t-test | | p = 0.0032 t=3.018, df=99 |
| SFig 2E[1] | Neck length in CA1 | Sham n=49, SDV n=52 | Unpaired t-test | | p = 0.0819 t=1.758, df=99 |

| Statistics of Suppl. Figure 3 | | | | | | |
|---|---|---|---|---|---|---|
| **Figure panels** | | **n** | **Statistical analysis** | | **F-value** | **p-value** |
| SFig 3A | LTP consolidation (30-40 min *vs* 50-60 min) | Sham n=8, SDV n=8 | 2-way ANOVA | interaction | $F_{(1, 14)}$ = 3.655 | p = 0.0766 |
| | | | | time | $F_{(1, 14)}$ = 4.631 | p = 0.0493 |
| | | | | groups | $F_{(1, 14)}$ = 8.609 | p = 0.0100 |

| Statistics of Suppl. Figure 4 | | | | | |
|---|---|---|---|---|---|
| **Figure panels** | | **n** | **Statistical analysis** | **F-value** | **p-value** |
| SFig 4A | α1 | Sham n=6, SDV n=7 | Unpaired t-test | | p = 0.3363 t=1.005, df=11 |
| | α2 | | | | p = 0.2614 t=1.184, df=11 |
| | α5 | | | | p = 0.9369 t=0.081, df=11 |
| | β1 | | | | p = 0.3267 t=1.027, df=11 |
| | β3 | | | | p = 0.4908 t=0.7128, df=11 |
| | γ2 | | | | p = 0.5871 t=0.5594, df=11 |
